# Relationship between cardiac cycle and the timing of actions during action execution and observation

**DOI:** 10.1101/585414

**Authors:** E. R. Palser, J. Glass, A. Fotopoulou, J. M. Kilner

## Abstract

Previous research suggests that there may be a relationship between the timing of motor events and phases of the cardiac cycle. However, this relationship has thus far only been researched using simple isolated movements such as key-presses in reaction-time tasks and only in a single subject acting alone. Other research has shown both movement and cardiac coordination among interacting individuals. Here, we investigated how the cardiac cycle relates to ongoing self-paced movements in both action execution and observation using a novel dyadic paradigm. We recorded electrocardiography (ECG) in 26 healthy adult subjects who formed 19 dyads (7 comprised of two subjects and 12 subjects paired with an experimenter). Each dyad contained an action executioner and observer as they performed a self-paced sequence of movements. We demonstrated that heartbeats are timed to movements during both action execution and observation. Specifically, movements were less likely to culminate synchronously with the heartbeat, around the time of the R-peak of the ECG. The same pattern was observed for action observation, with the observers’ heartbeats occurring off-phase with movement culmination. These findings demonstrate that there is coordination between an action executioner’s cardiac cycle and the timing of their movements, and that the same relationship is mirrored in an observer. This suggests that previous findings of interpersonal coordination may be caused by the mirroring of a phasic relationship between movement and the heart.

## Introduction

A growing body of research, comprising physiological and psychological investigations, is consistent with the hypothesis that the central nervous system has access to cardiac information, and uses this information to guide behavior. One such instance is the relationship between the cardiac cycle and the timing of movement. One of the earliest observations in this vein was that walking rate and heart rate typically show a one-to-one mapping^1,2^. It was proposed that the rhythm of the heart might represent a pacemaker, guiding the timing of movement.

Crucially, the effect of the heart on movement is bidirectional, with both effects of the cardiac cycle on movement and effects of movement on cardiac output observed. First, anticipatory cardiac slowing in the fore period of a reaction-time task while the subject prepares to move was reported^3,4^. Secondly, the motor response was also found to vary depending on when in the cardiac cycle it occurred ^5,6^, with faster reaction times during atrial contraction. A similar effect has recently been observed for movement inhibition, with faster responses to stop cues during systole^7^. Much of this previous physiological work on the relationship between the cardiac cycle and movement has employed simple prescribed movements within reaction-time and stop signal paradigms.

Cardiac-movement effects have been attributed to baroreceptors^8^. Located within the aortic arch, the carotid sinuses and the coronary arteries, these stretch receptors represent one of the key communication channels between the heart and the brain. Baroreceptors are active during systole, conveying information on the timing and strength of the heartbeat to the central nervous system via the vagal and glossopharyngeal nerves, but are quiescent during diastole when there is no such information to convey^9,10^. The heart’s rhythmicity is modified by humoral and neural factors and the heart period can be increased or decreased according to signals conveyed by vagal and sympathetic efferents^9–12^. There is therefore a closed loop communication reflex mediated by baroreceptors that allows the timing of the heart cycle to be controlled.

Interoceptive signals, particularly those originating from the heart, have been purported to play a role in social cognition^13,14^. Simulation theories of action understanding posit that in order to understand another’s behavior, we simulate their perceptual, motor and bodily states in ourselves^15^. As such, an action is understood based on the resonance of an observer’s motor system^16^. Such ideas have since been formalized by hierarchical predictive models, wherein it is argued that action observation results in an observer generating an internal model of how they would perform it^17^. Such models are proposed to contain not only exteroceptive and proprioceptive predictions^18^, but also interoceptive predictions about how such an action might be performed^19^. That is, a good model contains information pertaining not only to the expected position of the body in space, and afferent tactile feedback from the effector, but also the somatovisceral conditions that would be generated by the execution of that action. These predicted conditions can then be used to infer the causes of another’s behavior, based on the most likely causes of those conditions in the self. Including interoceptive information in predictive models may provide unique information about others’ affective states, not otherwise accessible through exteroceptive and prioprioceptive inference alone, due to the ubiquitous role of somatovisceral states in emotion and feeling states^20–23^.

Findings of resonance beyond the central nervous system suggest that this mirroring effect may be more widespread than previously thought, extending into the peripheral nervous system^28–31^. Both external stimuli such as observing someone perform an action, and internal stimuli, such as mentally rehearsing an action, can elicit autonomic system activity^32,33^. Watching an actor lift a weight or run on a treadmill produces changes in the respiration rate of an observer, such that it increased linearly with the treadmill speed^34^. These findings support the idea that bodily systems involved in interoception may also be candidates for resonance. The direct matching mechanism proposes a match between the conditions of action execution and observation^16^. Therefore, if this mirroring effect does extend to the interoceptive system, we would predict that any relationship between the cardiac cycle and movement timing found during action execution should also be found during action observation.

One consequence of resonance might be interpersonal coordination. Research in the field of social neuroscience has found both movement and cardiac coordination between interacting individuals. In the movement literature, interpersonal coordination may refer to either mimicry or synchronization^29^. A distinction is generally made based on the temporal relations of the interacting individuals, with mimicry reflecting non-periodic matched movements and synchronicity used to describe movements temporally matched in time^30,31^. Early studies reported that listeners to a speaker tend to move in time with the rhythms of a speaker’s speech^32,33^ and mirror the postures of the speaker^34^. In other interactions, the direction of the effect is less apparent, with partners mimicking each other’s postural sway^35^ and linguistic forms^36^. In a recent study, groups of subjects (numbering between four and five individuals) constructed models out of Lego blocks^37^. They either worked collectively (where subjects took turns to work on one construction project) or individually (where subjects worked on their own construction projects side by side). More stable shared heart rate dynamics were observed in collective trials than individual trials, increasing across the course of collective trials^37^, suggesting that convergence was tied to the social nature of the task. Indeed, heart rate entrainment has been suggested as a mechanism underlying emotional commonality in social interaction, facilitating a sense of community between individuals^38^, empathy^39,40^ and team performance^41–44^. In support of this, a study in which pairs of expert improvisers were asked to mirror each other’s movements found that periods of both increased kinematic and subjective togetherness were characterized by higher heart rates and greater inter-player correlation of heart rate^45^. Findings of interpersonal coordination and its influence on measures of social competence are compatible with the hypothesis that resonance supports action understanding.

The existing literature can largely be summarized as reflecting two broadly different methodological approaches to studying the relationship between the heart and movement. The first approach has studied the relationship between cardiac events and evoked movements such as those elicited by paradigms designed to gauge reaction time ^5,6^ and response inhibition^7^. These studies tested discrete, single movements. The second has taken the opposite approach and analyzed cardiac and movement parameters in a number of unconstrained self-paced environments, typically within a social context^37,38^.

The current study was designed to answer the previously neglected question of whether cardiac-motor timing and inter-individual resonance are linked, bridging the gap between the two previous approaches, by investigating cardiac-motor synchronicity in both action execution and observation. Naturalistic actions are more likely to comprise a series of goal-directed self-paced movements than discrete, cued movements. Therefore, in the present study, we sought to investigate the bidirectional relationship between the cardiac cycle and movement during a sequence of meaningful self-paced movements. Subjects were divided into dyads and took turns to perform and observe a sequence of semi-controlled movements under experimental conditions, while electrocardiography (ECG) was recorded from both (Figure 1). In brief, the task required both subjects within a dyad to memorize a sequence of six movement locations and then take it in turn to replicate the sequence. In this way, subjects both executed and observed the movements. The timing of movement was recorded using touch-sensitive pads.

**Figure 1:**
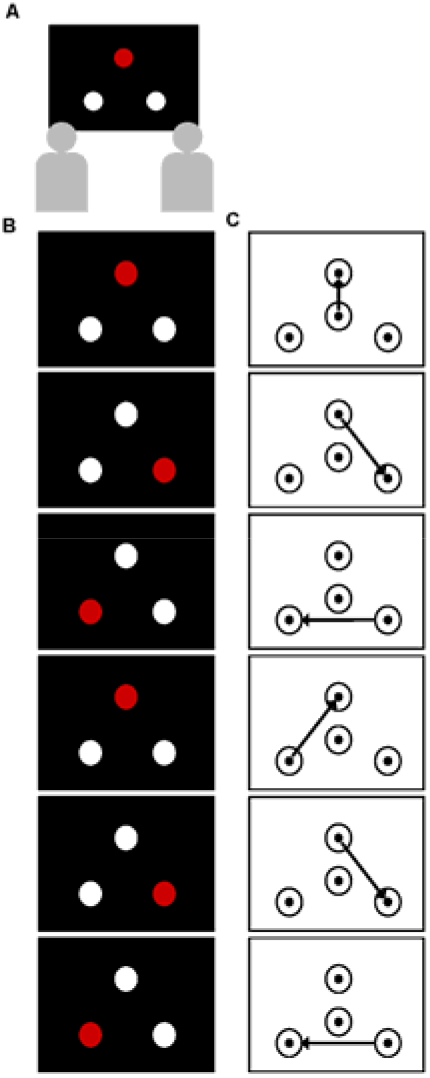
Task design. Panel A: Subjects were seated side by side in front of a laptop computer; Panel B: an example of the sequence of movements subjects were required to learn; Panel C: how they would accurately replicate the sequence in (B) using the touch-sensitive containers.

Firstly, we tested for coordination between execution and observation conditions within a dyad, in both heart rate and movement speed. Secondly, we investigated whether there was any relationship between the timing of the movements made, operationalized as the endpoint of the action, and the location of this event within the cardiac cycle, both in terms of its phase and time relative to R-peak. The endpoint of each action corresponded to each memorized location in the movement sequence. Thirdly, we tested the prediction that any phasic or time-based relationship observed between the timing of movement events and the cardiac cycle during action observation would also be observed during action observation.

## Results

Prior to testing our main hypothesis that there would be a relationship between movement timing and the period of the cardiac cycle, we tested for a significant difference in the length of the cardiac cycle (operationalized as the R-R interval) between observation and execution conditions. We also tested whether there was a relationship between the movement time of both subjects in a dyad and if there was a relationship between the length of the cardiac cycle and movement time.

There was no significant difference in the duration of the mean R-R interval between action execution and action observation (825.3 and 822.9 ms respectively; *t*(24) = 0.64, *p* = 0.53). There was no significant linear correlation between mean R-R interval and mean movement time. This was non-significant for both action execution (*r* = −0.37, *p* = 0.07) (Figure 2A) and action observation (*r* = −0.12, p = 0.58) (Figure 2B). However, there was a significant linear correlation between the movement time of subjects within a dyad (*r* = 0.61 *p* = 0.007 (Figure 2C). This significant relationship is consistent with the hypothesis that a common movement speed was adopted by the dyad but that this was not intrinsically tied to the length of the cardiac cycle.

**Figure 2:**
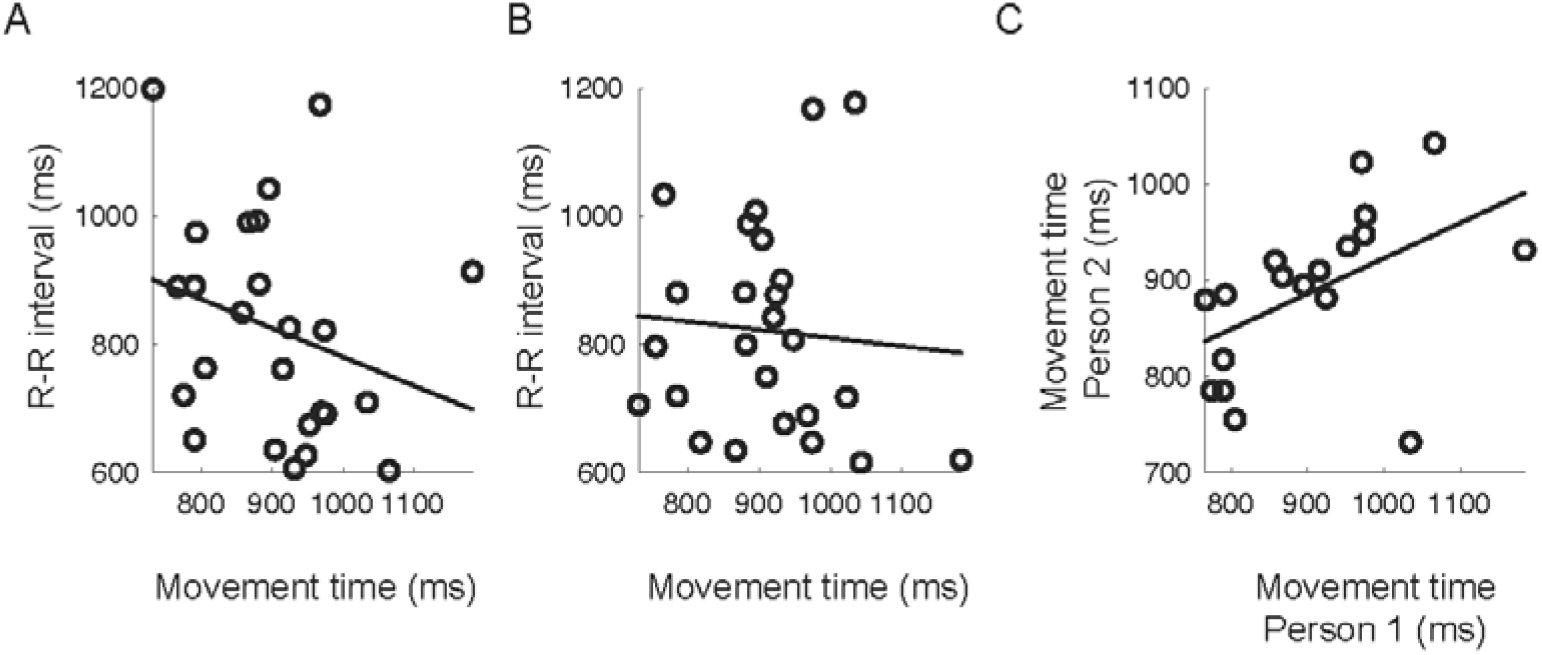
Relationship between cardiac and movement parameters. Panels A & B show the relationship between the mean movement time and the mean R-R interval for each subject, during action execution (A) and action observation (B). In each panel the circles show the data from each subject and the solid line shows the line of best fit. Panel C shows the relationship between the mean movement time for the two people that comprised the dyad. The significant positive relationship is consistent with the hypothesis that subjects imitated the movement speed of the other. In each panel, each circle shows the data from one dyad.

The main aim of this study was to test whether there was evidence of a consistent relationship between the timing of movement events and the cardiac cycle phase both when performing actions and when observing someone else perform them. This was tested both in the time and the phase domains.

### Time domain analysis

The data were divided into four time windows, centered around 0 (−100 – 100 ms), 200 (100 – 300 ms), 400 (300 – 500 ms) and 600 (500 – 700 ms) milliseconds post R-peak (Figure 3A). The repeated measures ANOVA with factors Condition (execution and observation) and Time (0, 200, 400 and 600) revealed a significant main effect of Time (*F*(2.21, 53.1) = 21.4, *p* < 0.001) and no significant effects of condition or interaction between the two factors (*F*(1,24) = 0.01, *p* = 0.92; *F*(2.39, 57.24) = 0.10, *p* = 0.93) respectively. Post hoc t-tests revealed that the significant effect of Time was driven by there being significantly less movement events in the 200 ms time period around the R-peak than the other three time periods (Figure 3B). For execution, the number of events in the −100–100 ms time window was significantly less than those in the 100–300 ms, and 300–500 ms time windows (Execution: *t*(24) = −4.59; *d* = −0.918, *p* < 0.05 & *t*(24) = −4.04, *d* = −0.81, *p* < 0.001). For observation the number of events in the −100–100 ms time window was significantly less than those in the 100–300 ms, 300–500 ms and 500–700 ms time windows *t*(24) = −7.404, *d* = −1.48, *p* < 0.05, *t*(24) = −4.91, *d* = −0.98, *p* < 0.001 & *t*(24) = −3.65, *d* = −0.73, *p* = 0.001). All other within Condition post hoc comparisons were non-significant when corrected for multiple comparisons.

**Figure 3:**
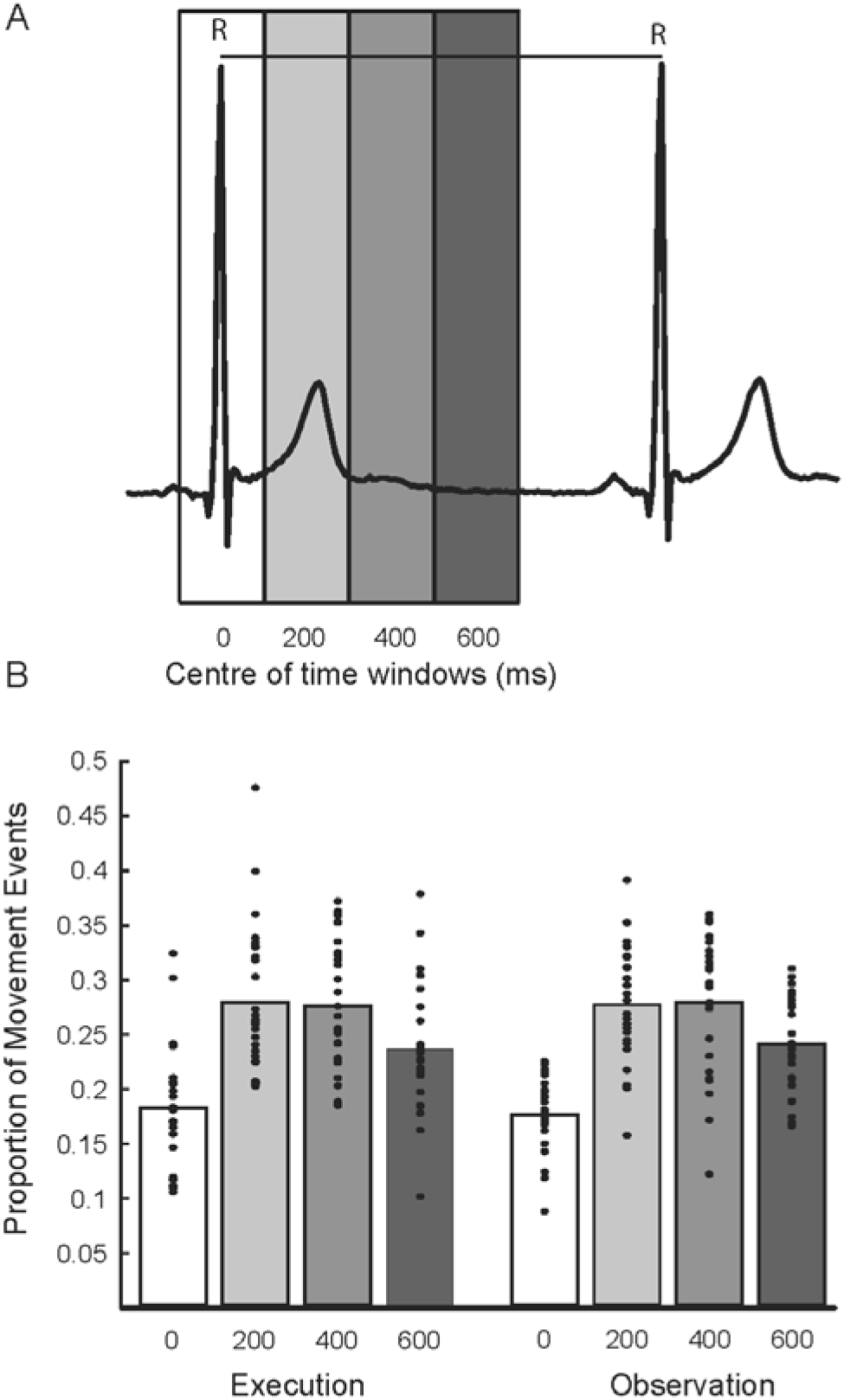
Relationship between time of movement culmination and cardiac cycle in the time domain. Panel A shows one cycle of the cardiac cycle (R-R) and shows the four time windows used for the analysis. Panel B shows the mean proportion of events that occurred in each of the four windows shown in A. The dots show the mean data for each subject, the bar is the mean across subjects.

### Phase domain analysis

In contrast to the time-domain analysis, there was no evidence that the mean phase of each subject’s movements differed significantly from the uniform distribution in either the Execution or Observation conditions (Rayleigh test *z* = 0.07, *p* = 0.93 & *z* = 0.84, *p* = 0.43 respectively (Figure 4A & B)). To investigate the phase data further we analyzed the mean distribution of the phase of all events across subjects during action execution and observation conditions. If there was no consistent phase relationship these values should be close to the excepted value for a uniform distribution (Figure 4C & D). Here, with 20 phase bins, the proportion in each bin with a uniform distribution would be 0.05 (1/20). However, in both conditions there were more motor events than expected at particular phases of the cardiac cycle (Figure 4C–F). For the Execution condition there were significantly more events than expected in the phase bin centred on 81° (*t*(24) = 2.07, *p* = 0.049). For the Observation condition this occurred later in the cycle, in the phase bin centred on 153° (*t*(24) = 2.32, *p* = 0.029). However, it should be noted that these were small effects that did not survive correction for multiple comparisons. There was a reduction in the proportion of events that occurred either before and slightly after the R-peak during Execution, or after the R-peak during Observation. However, these reductions were not significantly less than expected.

**Figure 4:**
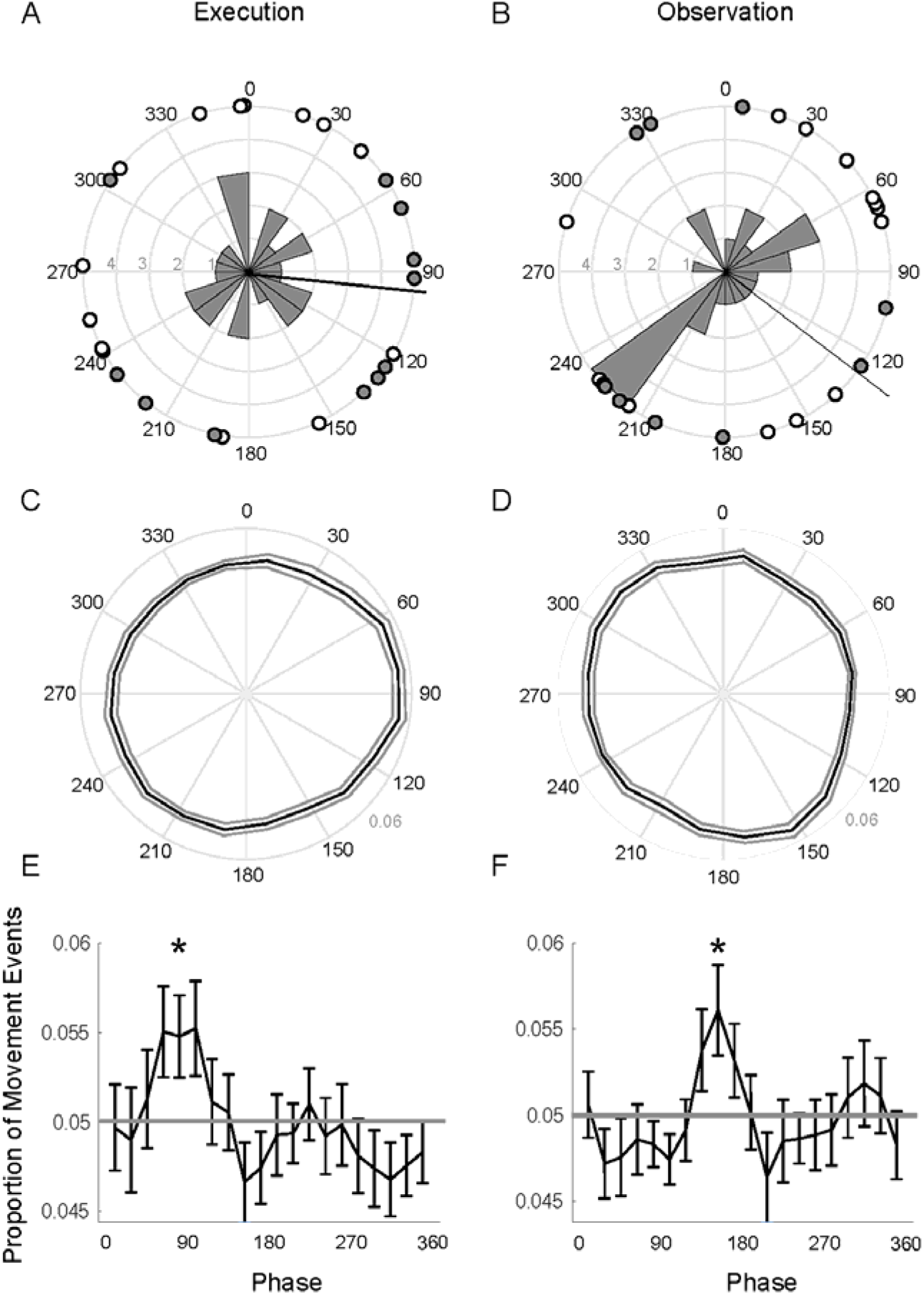
Relationship between time of movement culmination and cardiac cycle in the phase domain. Panels A-F show the relationship between the time of the movement event (action end point) with respect to the phase of the R-R interval. In each panel the R peaks occurred at 0° and the phase through the R-R interval cycles clockwise around the circle. On the left-hand side of figure, panels A, C & E show the data for executed actions and on the right-hand side, panels B, D & F show the data for observed actions. In A & B the circles show the mean phase for each subject. White circles are from those dyads where the ECG was recorded from both subjects and the grey circles show the data where ECG was recorded from only one subject. The circular histogram show the frequency of the data as a function of phase. The solid black arrow shows the circular mean phase across subjects. Panels C & D depict the distribution of the mean normalized circular histograms across subjects. The solid black line depicts the mean phase distribution and the dotted lines show the standard error around this mean (SEM). Panels E & F show the same data in C & D but unwrapped. The solid grey line shows the null distribution (a uniform distribution). Error bars represent the SEM. * show phases where the proportion of events are significantly different from a uniform distribution.

## Discussion

Here, we showed that the timing of self-paced movements and phases of the cardiac cycle are linked in a similar way for both action execution and observation. We showed that movement events were significantly less likely to occur at the time of the heartbeat, synchronous with the R-peak, and significantly more likely to occur between heartbeats. This same relationship was also observed between the motor events of an actor and the cardiac cycle of an observer. That is, the observer was significantly less likely to experience a heartbeat when observing movement endpoints. Additionally, there was a significant positive correlation between movement time in execution and observation conditions, suggesting dyads adopted a common movement speed. These results build on previously separate literatures that find both a relationship between discrete motor events and cardiac timing, and cardiac and movement coordination between interacting individuals.

In this study we investigated the relationship between movements and the cardiac cycle in both the time and phase domains. The effect of a relationship between the two was largest in the time domain analysis. Indeed, despite there being a clear and significant reduction in the proportion of the events around the time of the R-peak in the time domain analysis the group level phase analysis showed no evidence that the distribution was non-uniform. The time domain analysis assumes that any cardiac effects are time locked to the R-peak and are independent of the R-R interval. Whereas the phase analysis assumes that any cardiac effects are locked to a particular phase of the R-R interval and are not time locked. This difference in the effect sizes reported here most likely reflects these different sensitivities of the two analyses. Future work needs to focus on studying the different specificities and sensitivities of the different analysis pathways to understand when they should be used and how to understand the differences between them.

Many have suggested that an internal central pacemaker guides the timing of motor actions^1,2,49,50^. The heart, with its regular rhythmicity that modulates with arousal level, intuitively represents a good candidate for this role. Information on the strength and timing of heartbeats is conveyed to the central nervous system via the vagal and glossopharyngeal nerves. Such information could thus be capitalized on by the motor system when timing the occurrence of actions. Indeed, there is evidence that stimulation of afferent vagal and glossopharyngeal pathways is associated with modulations of efferent motor pathways. For example, pressure applied to the carotid sinus, which modulates baroreceptor firing^51^, inhibits spontaneous movements and reduces muscle tone in anaesthetized animals^52^. Efferent projections, which can delay the onset of the next heartbeat, could further optimize control over timing relationships.

The benefits of timing behavior to the cardiac cycle, however, remain unknown. Global perceptual attenuation during heartbeats has been proposed, as sensations are inferred as consequences of internal cardiac activity and therefore down-weighted^53^. Indeed, a recent study purports that exteroceptive sensory confidence is greatest during diastole and at its lowest concomitant with the heartbeat^54^, providing a neat explanation for the preference to time movements in between heartbeats observed here.

It is possible that actions are timed to the point in the cardiac cycle where the ability to balance autonomic and behavioural demands is greatest. Previous research has shown that as an individual attends to and prepares to interact with a stimulus, vagal activation lengthens the cardiac cycle^55^. In turn, vagal withdrawal, when the individual initiates a response, shortens the cardiac cycle^56,57^. However, the timing of the behavioral response also impacts the cardiac consequences. Responses made at or just after the R-wave impact the length of the current cardiac cycle, whereas responses made later in the R-R interval impact the length of subsequent cycles. Furthermore, the sensitivity of the sinoatrial node to input from the vagus is not uniform across the cardiac cycle. Instead, a period of heightened sensitivity is estimated to occur between 250 and 400 milliseconds post R-wave, followed by a period of relative insensitivity^58,59^. This sensitive period broadly corresponds with the period of increased movement events in the current study. Considering that sympathetic efferents typically impact heart rate relatively slowly, on a time–scale of between 1,300 and 2000 milliseconds^60,61,62^, timing movement events may capitalize on these effects to more rapidly adjust autonomic outflow to match demand.

Additionally, it is possible that as agents, we act upon the environment such that relevant signals appear during optimal periods of the cardiac cycle^63^. It may be that stimuli whose processing would benefit from information on the state of cardiovascular arousal are more acutely perceived at systole, when baroreceptor feedback occurs, while stimuli that would not benefit from such signalling are better processed during diastole. As such, during systole, fearful faces are more readily detected, and result in greater amygdala activation^64^, and race-related threat appraisal is heightened^65^. Experimentally increasing heartrate (and therefore the amount of time spent in systole) heightens fear responses but not disgust responses^66^. Meanwhile, during diastole, target shooting^67, 68^ and memory for words^69^ is more accurate. The current results suggest we may act upon our environment in such a way as to maximize this interoceptive distinction.

Respiration was not measured in the current study, but could play a role in the current findings. A recent study identified a correspondence between the timing of voluntary movement and the respiratory cycle, with movement initiated more frequently during expiration^70^. Expiration coincides with a decrease in heart rate, compared to the increase observed during inspiration. Therefore, movements made during inspiration are more likely to coincide with heartbeats than those made during expiration. We have greater voluntary control over respiratory than cardiac events, control that could be leveraged to better govern cardiac timing. Holding one’s breath or breathing out slowly when performing a precise movement may be one way of decreasing the chances of a heartbeat occurring at the crucial moment.

In the present study, we not only observed a relationship between the cardiac cycle and action execution, but also observed a similar relationship for action observation. Observation of another may produce resonance of the mechanisms involved in executing that action. Activation may remain sub-threshold for actually producing the motor output, however associated somotovisceral changes may be produced^28^. Afferent interoceptive feedback resulting from these changes may in turn support inferences about the causes of the other’s behavior. As such, it is possible that inference about the mental states of others is supported by a form of interoceptive resonance or mirroring. Such a mechanism might explain how, in the present data, action observation resulted in a similar movement-cardiac rhythm to that seen in the individual executing the action. As the observer’s motor system predicts the actor’s next movements, comparable interoceptive conditions are simulated, much like action observation results in an increase in excitability in the muscles required to perform the observed action^15^.

The mutual prediction of each other’s’ actions is thought to facilitate joint coordinated action and the achievement of shared goals^71–73^. There is evidence to suggest that shared representations of action form automatically even when no joint action is required and it would be more effective to ignore the other person^74^. The results found here suggest the possibility that cardiac-movement synchronicity may support the interpersonal coordination that underlies joint action, through physiological attunement. Such coordination may serve as “social glue”, fostering feelings of group belonging and closeness^75^. A number of findings appear to support this. Professional interviewers’ judgements of trainees’ competence have been found to correlate positively with the trainees’ convergence of speech rate and response latency^76^. The degree of convergence appears to vary based on personality traits and social standing. For example, subjects who scored higher on a measure of the need for social approval converged more to their partner’s speech intensity and pause length than subjects who scored lower^77^. Moreover, greater language convergence is observed towards occupational superiors than subordinates, in that foremen converge more to the language of managers. Increased physiological covariation, in the form of galvanic skin response (GSR) has been observed in dyads who either like or dislike each other, compared to those who hold neutral opinions of each other^78^. Future work should seek to investigate whether inter-dyad factors such as inter-individual liking and physical similarity to one’s partner, or individual differences such as personality traits, are predictive of the cardiac-movement coordination observed here.

Previous investigations have suggested that the influence of interoceptive signals on behavior may be mediated by the individual’s degree of conscious awareness of those signals^69,79^. However, it is unclear whether the present effects rely on conscious control of behavior, or reflexive processes below the level of conscious awareness. In a previous study investigating the effect of the cardiac cycle on timing preferences for actively sampling the environment, operationalized as the time subjects chose to press a key prompting a visual stimulus relative to their cardiac phase, no effect of inter-individual differences in awareness of interoceptive signals was observed^63^. If the action observation effects observed here are consciously mediated, it raises the possibility that a greater awareness of cardiac signals might facilitate greater social mimicry. Future research could test the possibility that inter-individual variability in the action observation effect reported here is related to social competencies and is mediated by interoceptive awareness.

Important questions remain regarding precisely under what conditions cardiac-movement coordination occurs. Understanding this may aid in interpreting previous findings that report no relationship between the heart and movement^80^. At the level of a single subject, it is likely that not all centrally mediated sensorimotor events show a phased relationship with the heart. It is perhaps noteworthy that in the present study, many of the subjects reported during debriefing that they found replicating the sequence of movements challenging. The cognitive demands of the task may be an essential difference between the present design and other studies that report no relationship between heart and movement. In dyads, possible mediators include the history of interpersonal interaction, speech coordination, the structure of the task and the dynamics of turn-taking, and the degree of behavioral coordination between the agents^39^. Different patterns of kinematic and autonomic coordination have been reported. Under some circumstances, a leader-follower pattern is observed^32,34^, while in other scenarios, convergence appears to be more bi-directional^35–37^. It would be interesting for future research to explore how the heart rates and kinematics of individuals in a dyad change over time. A plausible prediction is that over the course of a trial, the observer’s heartbeats become increasingly linked to the timing of the actor’s movements. Finally, research suggests an association between elevated blood pressure and reduced social competence^81,82^. As baroreceptor activity is tied to blood pressure, the current findings raise the possibility that reduced cardiac mirroring may underlie these findings.

In summary, these findings demonstrate a relationship between the timing of motor events and the cardiac cycle in both action execution and observation. These results suggest that interoceptive signals from the heart and self-paced movement are intrinsically linked. The observation of this phasic relationship both when executing and observing action raises the possibility that this relationship may be driving interpersonal coordination effects, which have been found to foster successful social interaction.

## Materials and Methods

### Subjects

A total of twenty-six healthy adult subjects with normal or corrected-to-normal vision were recruited. Subjects were aged between 19 and 36 years (mean = 24.92, *SD* = 4.78). Fourteen were female and 12 were male. Twenty-four subjects reported being right-handed, and two reported being left-handed. Data was collected in two stages. In the first stage, due to practical reasons, subjects performed the dyadic behavioural turn-taking task with an experimenter as their partner (n = 12). In stage two, the remaining subjects were recruited in pairs and performed the task together (n = 14). For analysis, data were pooled (see Supplementary Materials for the results reported separately). Ethical approval of the experimental protocol was granted by the Ethics Committee of University College London, and the methods were carried out in accordance with the Declaration of Helsinki. All subjects gave written informed consent before testing.

### Materials

Subjects were seated in pairs at a table in front of a Dell laptop computer. Stimuli were presented using MATLAB (version 7.8.0, Mathworks Inc., MA, USA) with the Cogent 2000 toolbox (www.vislab.ucl.ac.uk/cogent_2000.php). Three electrodes were affixed to the abdomens of subjects as per the standard Lead II configuration, to record their ECG. On the table were four touch-sensitive pads and a marble, which the subjects used to execute a series of memorized movements.

### Procedure

Once seated comfortably, the experimenter affixed three disposable or washable electrodes to each subject. They were instructed that they would view movement sequences on the screen (see Figure 1A) and would have to replicate them in turns afterwards using the marble and touch-sensitive pads in front of them. They were also instructed not to talk to or interact with their partner for the duration of the experiment. On the screen, subjects viewed an animation of a sequence of six movements (see Figure 1B). The screen showed three white circles arranged in a triangle. One circle briefly turned red before returning to white to indicate the sequence. The order of the sequence was pseudo-randomized with the only constraint being that the same circle could not appear two or more times in a row. Subjects viewed the sequence twice and were instructed to memorize it. Subjects were informed that the ECG measures their heart, but were not made aware of the purpose of the experiment or that the timing of their movements was being recorded.

On the table were four touch-sensitive pads and a marble. Three of the pads were arranged in a triangle and corresponded to the circles that appeared on the screen. The fourth pad was placed in the middle and was the home pad where the marble was to be returned to at the end of each movement sequence and from where each movement sequence commenced. On each trial both subjects in the pair had to attempt to replicate the sequence, one after the other. To do this, subjects were instructed to move the marble between the touch sensitive pads in the order they believed to be the same as in the initial video (see Figure 1C). Subjects were instructed to make the series of movements at a speed that was comfortable to them.

The order of which subject executed first and which second was counter-balanced across blocks for each pair. That is, turn-taking occurred by block, where the same subject took the first turn for a whole block, and then they swapped. Which subject was instructed to go first was alternated, so that no subject took the first turn for more than one block in a row. For 12 subjects there were two blocks of ten trials in total. In one block one subject would go first for all ten trials, and in the other the other subject would go first for all ten trials. For the remaining 14 subjects there were four blocks of ten trials in total. In two blocks one subject would go first for all ten trials, and in the other two the other subject would go first for all ten trials. ECGs were recorded throughout the duration of each block.

For 18 subjects the ECG was recorded using an Active 2 Biosemi amplifier. The ECG was recorded at 2048 Hz. For the remaining seven subjects, the ECG was recorded using a CED 1401 in Spike2 at a sampling rate of 1000 Hz. The time at which the marble was placed on the touch-sensitive pad was also recorded. The experiment lasted approximately 40 minutes. Afterwards, subjects were debriefed about the purpose of the experiment and their experience of taking part, and were given the opportunity to ask the experimenter any questions.

### Analysis

For all subjects the three electrode ECG recordings were transformed offline by linear subtraction into two sets of bipolar recordings, which in turn were averaged to produce a single ECG recording for each subject. For all subjects, the time of the peak of the R-wave of each heartbeat was calculated from the ECG. First, the ECG data were high pass filtered at 0.01Hz to remove any linear drift. Second, a threshold was determined from the data to isolate the R-wave and the time point of local maximum for each suprathreshold peak was calculated. For each subject, this time point was determined at 1000 Hz (i.e., to millisecond accuracy) and in this way all data were then encoded at the same sampling rate. For each block the time the touch-sensitive pads were contacted was extracted from the event channel and also resampled at 1000 Hz where necessary. For one subject there was a failure of the touch-sensitive pads and this subject was excluded from further analysis, leaving 25 subjects for data analysis.

### Analysis of movements and phase of the cardiac cycle

To address whether or not there was a statistical relationship between the time the marble was placed on the touch-sensitive pad and the phase of cardiac cycle two different analyses were performed, one in the time domain and one in the phase domain.

### Time domain analysis

In the time domain the proportion of motor events that occurred in four time windows relative to the R-peak were calculated. The four time windows were −100–100 ms, 100 –300 ms, 300 –500 ms and 500 –700 ms, that is, centered around 0, 200, 400 and 600 ms respectively (Figure 3A). The proportion of motor events that occurred in these windows was calculated for both executed and observed events. To determine if there was any effect of time on the proportion of events that occurred a repeated measures ANOVA was run with two factors, Condition (execution and observation) and Time 0, 200, 400, & 600). Significant effects were further investigated with post-hoc t-tests, Bonferroni corrected for multiple comparisons).

### Phase domain analysis

In the phase domain, we calculated the phase of the movement event as a function of the R-R interval^65^. Circular statistics were employed in order to exploit the repeating nature of the cardiac cycle. To this end, for each action endpoint the time of both the preceding and proceeding R-wave was calculated and the phase of the action endpoint was calculated as a function of the R-R interval in which it occurred. For example, for an R-R interval of time t_R,_ where the movement event occurred at time t_e_ the phase, in degrees, of that event was calculated as t_e_/T_R_ × 360. For each subject, the mean phase was then calculated for the execution and observation blocks separately using circular averaging. This resulted in two mean phases per subject, one for each condition (execution and observation). We then tested, separately for each condition, whether these phases differed from uniformity using Rayleigh tests. In addition, we also calculated the distribution of phases in a circular histogram, where each bin was a 20^th^ of a circle. These data were normalized so each bin represented the proportion of total events for each subject for each condition. These were smoothed by averaging over three neighbouring bins, the central bin and the two neighbouring bins. The first analysis shows the uniformity of the mean phase across subjects but gives no indication of the degree of uniformity of the phases for each subject. This is shown by the second analysis.

## Acknowledgements

This work was supported by a PhD studentship awarded to E. R. Palser from the Economic and Social Research Council (ESRC) and the Medical Research Council (MRC). A. Fotopoulou was supported by a ‘European Research Council Starting Investigator Award’ [ERC-2012-STG GA313755].

## Author Contributions

Designed the research ERP, AF, JMK; collected the data ERP, JG; analyzed the data ERP, JMK; wrote the manuscript ERP, JMK; provided comments on a draft and approved the final manuscript AF, JG.

## Competing interests

The authors declare that they have no competing interests

## Notes

### Competing Interest Statement

The authors have declared no competing interest.

