## Supplementary Materials for "Relationship between cardiac cycle and the timing of actions during action execution and observation"

Data collected in stage one and two were pooled for statistical analysis, however we show here that comparable results were observed in both groups when analyzed separately.

*Time domain analysis*

Stage 1: The repeated measures ANOVA with factors Condition (execution and observation) and Time (0, 200, 400 and 600) revealed a significant main effect of Time ($F(2.11, 21.11) = 10.11$, $p = 0.001$) and no significant effects of condition or interaction between the two factors ($F(1,10) = 1.31$, $p = 0.28$; $F(2.19, 21.85) = 0.24$, $p = 0.808$) respectively.

Stage 2: The repeated measures ANOVA with factors Condition (execution and observation) and Time (0, 200, 400 and 600) revealed a significant main effect of Time ($F(1.95, 25.30) = 13.28$, $p < 0.001$) and no significant effects of condition or interaction between the two factors ($F(1,13) = 0.26$, $p = 0.62$; $F(2.43, 31.63) = 0.53$, $p = 0.63$) respectively.

*Phase domain analysis*

Supplementary Figure 1: Here we show the t-statistic at each phase for the three populations: Stage1 (green), Stage 2 (red) and All subjects (blue). Panel A shows the data for execution and Panel B for Observation. The * show phases where the effects were significant. As is clear both groups show a qualitatively similar pattern of modulation with phase.

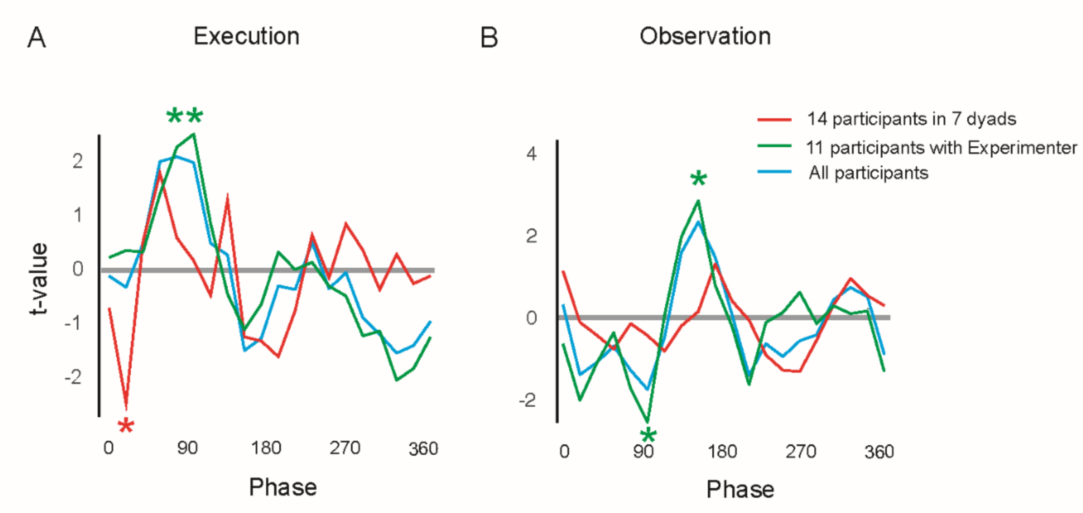
A
Execution
B
Observation
14 participants in 7 dyads
11 participants with Experimenter
All participants
t-value
Phase
Phase
0
90
180
270
360
